# Short-term plasticity in the visual thalamus

**DOI:** 10.1101/2021.10.14.464354

**Authors:** Jan W. Kurzawski, Claudia Lunghi, Laura Biagi, Michela Tosetti, Maria Concetta Morrone, Paola Binda

## Abstract

While there is evidence that the visual cortex retains a potential for plasticity in adulthood, less is known about the subcortical stages of visual processing. Here we asked whether short-term ocular dominance plasticity affects the visual thalamus. We addressed this question in normally sighted adult humans, using ultra-high field (7T) magnetic resonance imaging combined with the paradigm of short-term monocular deprivation. With this approach, we previously demonstrated transient shifts of perceptual eye dominance and ocular dominance in visual cortex (Binda et al., 2018). Here we report evidence for short-term plasticity in the ventral division of the pulvinar (vPulv), where the deprived eye representation was enhanced over the non-deprived eye. This pulvinar plasticity effect was similar as previously seen in visual cortex and it was correlated with the ocular dominance shift measured behaviorally. In contrast, there was no short-term plasticity effect in Lateral Geniculate Nucleus (LGN), where results were reliably different from vPulv, despite their spatial proximity. We conclude that the visual thalamus retains potential for short-term plasticity in adulthood; the plasticity effect differs across thalamic subregions, possibly reflecting differences in their cortical connectivity.

## INTRODUCTION

A classic paradigm for probing brain plasticity is monocular deprivation. During development, patching one eye for several days weakens the cortical representation of the deprived eye producing a stable change of ocular dominance columns in primary visual cortex (Hensch & Quinlan, 2018; Wiesel & Hubel, 1963, 1965). In adult humans, a much shorter period of eye patching (about two hours) produces a paradoxical enhancement of the deprived eye signal (Bai et al., 2017; Binda & Lunghi, 2017; Castaldi et al., 2020; Chadnova et al., 2017; Lunghi, Berchicci, et al., 2015; Lunghi et al., 2011; Lunghi et al., 2013; Lunghi, Emir, et al., 2015; Lunghi & Sale, 2015; Lunghi et al., 2019; Lyu et al., 2020; Min et al., 2018; Schwenk et al., 2020; Wang et al., 2020; Zhou et al., 2015; Zhou et al., 2013; Zhou et al., 2014) that was interpreted as a form of homeostatic plasticity (Turrigiano, 2012). Recently, we explored the neural underpinnings of this effect using ultra-high field functional Magnetic Resonance Imaging (fMRI). Although our technique did not directly measure ocular dominance columns, we were able to detect short-term plasticity effects in primary visual cortex V1 that were compatible with a change in ocular drive (Binda et al., 2018).

While ocular dominance plasticity has been thoroughly investigated in the visual cortex, less is known about its effects on subcortical visual processing. The thalamus is a crucial node of the visual system, with different subnuclei serving complex and diverse functions that we have only begun to uncover. The Lateral Geniculate Nucleus (LGN) receives the largest contingent of retinofugal fibers, and it is the main source of feedforward signals to V1 (Blasdel & Lund, 1983; Hendrickson et al., 1978; Hubel & Wiesel, 1972). In humans, there are indications that LGN can shift function following sensory deprivation (Levine et al., 2020; Mikellidou et al., 2019) or restoration (Castaldi et al., 2016). In rodents, plasticity of lateral geniculate nucleus neurons was recently reported (Jaepel et al., 2017; Rose & Bonhoeffer, 2018; Sommeijer et al., 2017). These data open the possibility that short-term ocular dominance plasticity originates in the thalamus and it is projected via feedforward connections to the visual cortex of human adults.

Adjacent to LGN, the Pulvinar is the largest thalamic nucleus that processes visual information; it has been repeatedly implicated in developmental plasticity of the visual system (Bourne & Morrone, 2017; Bridge et al., 2016). Although it receives signals from the retina and the superior colliculus, the primary input to Pulvinar comes from a range of cortical areas, with different subnuclei showing distinct connectivity patterns. Cortical connections are typically bidirectional, supporting cortico-thalamo-cortical loops for information exchange (Saalmann & Kastner, 2011). The function of these loops is only partially understood; they may help control response gain in the cortex, effectively regulating the balance between competing visual signals (Fiebelkorn & Kastner, 2019; Purushothaman et al., 2012) As such, they may be ideally positioned to regulate eye-specific response gain and thereby mediate ocular dominance plasticity effect.

Thus, in principle, both LGN and ventral Pulvinar may support plasticity. However, no previous study has tested their potential for short-term reorganization in the human adult. Here we address this question using the short-term monocular deprivation (MD) paradigm applied to normally sighted human adults studied with Magnetic Resonance Imaging (MRI). Mapping thalamic nuclei with MRI is notoriously difficult due to the low signal to noise ratio and the small size of these structures. Here we overcome these limitations using ultra-high field (7T) fMRI and relying on independently defined regions of interest (from the Natural Scenes Dataset, Allen et al., 2021). These label LGN and two subdivisions of the Pulvinar, ventral (vPulv) and dorsal. vPulv is most tightly connected with occipito-temporal visual cortex and it is retinotopically organized; the ROI comprises two complete and precise maps of the contralateral hemifield, sharing the fovea and the vertical meridian; it partially overlaps the anatomically defined lateral and inferior pulvinar subnuclei. Conversely, the dorsal region of the Pulvinar shows rough retinotopy and its responses are better explained by attentional and cognitive phenomena than by the low-level visual features, which is coherent with its preferential connectivity with parietal cortex (Arcaro et al., 2018; Arcaro et al., 2015). Thus, vPulv is most clearly involved in visual processing and should respond most vigorously to the simple visual stimuli used in our study; for these reasons, we focused our analyses on this region and compared its behavior with LGN.

## RESULTS

We measured 7T BOLD responses to monocular visual stimulation delivered before and after 2h of eye patching in 18 adult participants with normal vision (Experimental design is shown in **Figure 1A**). We previously analyzed responses in visual cortical areas (Binda et al., 2018); here we analyzed responses in the visual thalamus. Pooling data across participants, after aligning them to the MNI template (Avants et al., 2008), we found that visual responses within the thalamus clustered in two foci (**Figure 1B**) that match two thalamic regions with established visual properties (colored outlines in **Figure 1B**): Lateral Geniculate Nucleus (LGN) and ventral Pulvinar (vPulv), localized based on independent work (they were obtained from the Natural Scenes Dataset NSD; Allen et al., 2021).

**Figure 1.**
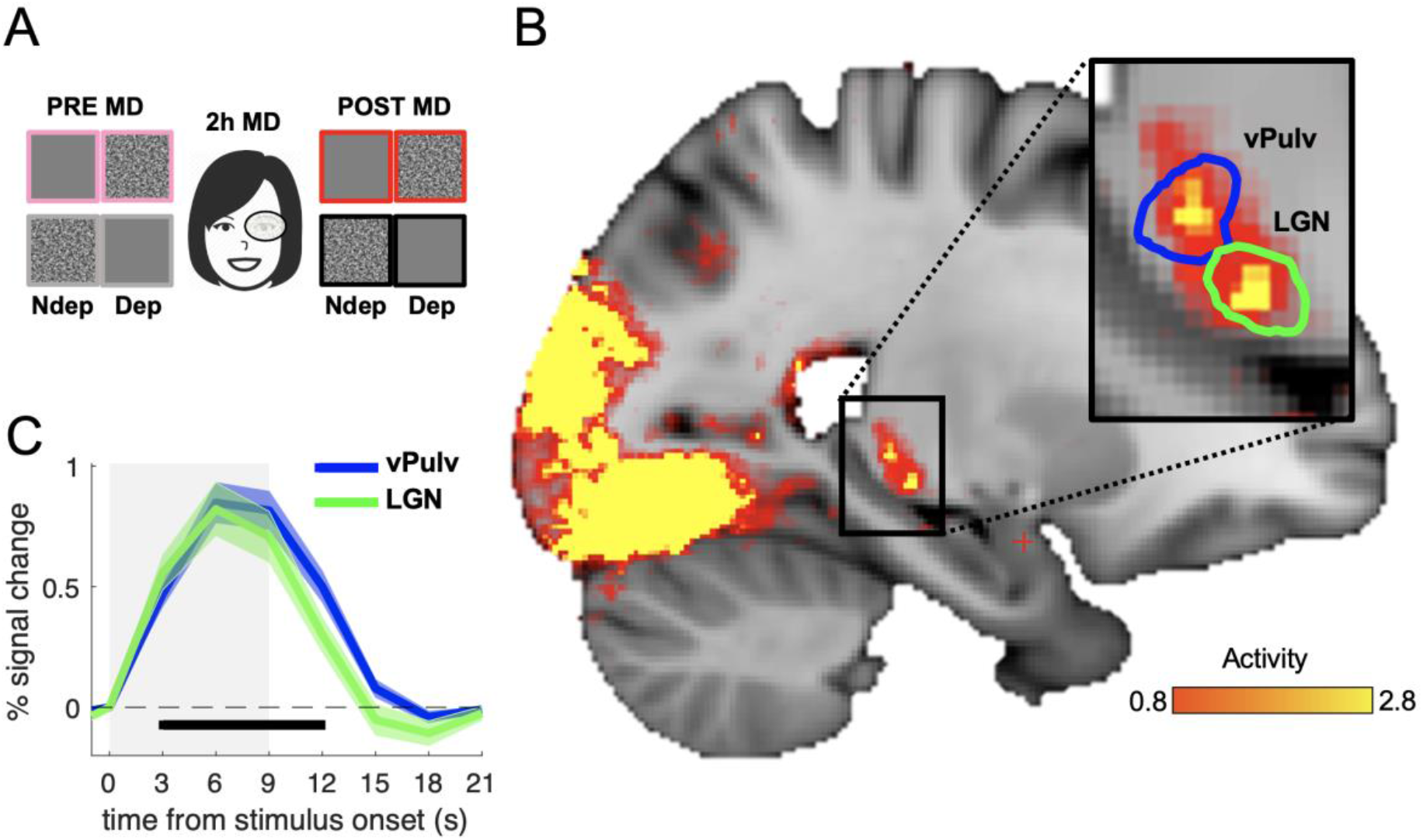
Visual responses in the thalamus. **A)** Experimental design. Responses to monocular presentations of dynamic noise filtered at various spatial frequencies were recorded before and after monocular deprivation. Binocular rivalry was measured immediately before each scanning session and used to estimate perceptual eye dominance. **B)** Map of visually evoked activity, estimated by fast Fourier analysis of the BOLD time series (amplitude at stimulus frequency divided by the root mean squared error of the corresponding sinusoidal function; data pooled across all sessions, stimulating both eyes before and after deprivation). Colored lines outline the two independently defined subcortical ROIs (Allen et al., 2021) where we focused our analyses: ventral pulvinar (vPulv) and Lateral Geniculate Nucleus (LGN). **C)** Temporal dynamics of the BOLD response (gray shaded area represents stimulus duration) in the two subcortical ROIs; curves and surrounding shaded areas show means and standard errors across participants (data pooled across all sessions and averaged after subtracting the baseline BOLD signal at stimulus onset). The black bar marks timepoints where the BOLD response in both ROIs is above zero, significant at p < 0.05.

**Figure 1C** shows the temporal dynamics of BOLD responses (to stimuli in the to-be-deprived eye, before monocular deprivation) extracted from these independently defined ROIs. Responses had similar size in LGN and vPulv. These were clearly weaker than previously measured in V1 (were signals peaked at about 2.5% at 9s from stimulus onset; Binda et al., 2018), but reliably larger than 0 at all points between 3s after stimulus onset to 3s after its offset (all t(17) > 4.30 and p < 0.01). Response dynamics was faster than in V1 (the peak here occurs around 6s from stimulus onset), and slightly faster in LGN than in vPulv, as previously reported (Lewis et al., 2018). This led us to quantify BOLD response amplitudes with an approach that makes minimal assumptions on temporal dynamics. Since the visual stimulus was a periodic alternation of stimulus present/absent epochs (ignoring variations in a stimulus dimension that is not relevant here, see methods), the amplitude of visually evoked responses could be extracted simply by Fourier analyses of the fMRI timeseries, taking the amplitude at the stimulus frequency (note that analyses based on General Linear Modelling and Event Related Averaging produced the same pattern of results, as detailed below).

With this approach, we compared responses to stimuli delivered to the two eyes.

Before monocular deprivation, no systematic differences in eye dominance were expected; therefore, we used BOLD responses to stimuli in the two eyes for estimating the internal consistency of our results. We found that responses to the two eyes were correlated across participants in both thalamic regions (Pearson’s correlation coefficients were *r*(*18*) = 0.66, p = 0.003 for LGN and *r*(*18*) = 0.58, p = 0.011 for vPulv), indicating good test-retest reliability of our measurements and allowing us to examine their variations after monocular deprivation.

**Figure 2** compares responses before and after deprivation to stimuli in the deprived and non-deprived eye. vPulv showed a significant eye by time interaction (**Figure 2A**: F(1,17) = 14.75, p = 0.001), similar as that seen in V1 (Binda et al., 2018). This is the hallmark of a significant short-term plasticity effect. In contrast, LGN showed no significant effect (**Figure 2B**, F(1,17) = 0.18, p = 0.675). This indicates that monocular deprivation selectively boosted responses to the deprived eye in the ventral Pulvinar, but it did not reliably affect the Lateral Geniculate Nucleus. The three-way interaction of factors eye, time and ROI was significant (F(1,17) = 5.76, p = 0.028), implying that the two thalamic regions were systematically different in their response to monocular deprivation and suggesting that noise in LGN responses could not account for the lack of plasticity effect in this ROI. As previously seen in V1 (Binda et al., 2018), we found that the inter-individual variability of the plasticity effect-size in vPulv was physiologically meaningful, as it correlated with the size of the behavioral effect (**Figure 2C**; *r*(*18*) = 0.47, p = 0.048); on the contrary, no significant correlation was found for the effect in LGN (*r*(*18*) = −0.12, p = 0.645, not shown).

**Figure 2.**
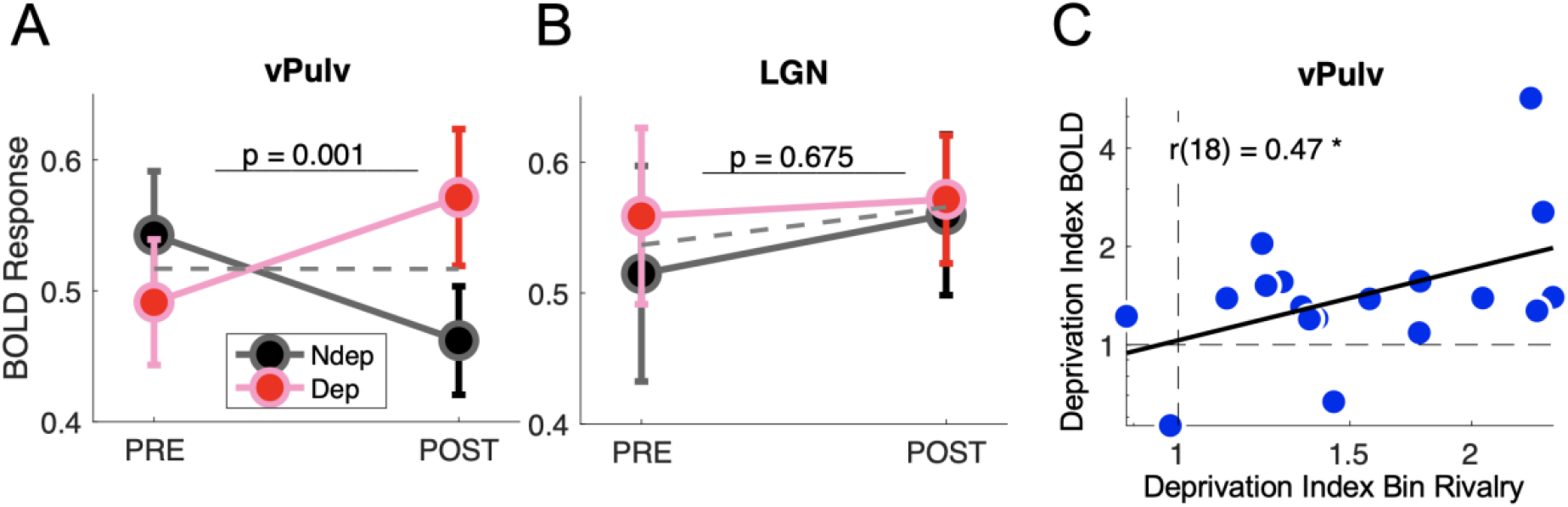
Short-term plasticity in Pulvinar, not in LGN. **A-B)** Modulation of visually evoked BOLD responses with monocular deprivation, in the deprived and non-deprived eye. BOLD responses were quantified by fast Fourier analysis of the fMRI timeseries, taking the amplitude at the stimulus frequency. Symbols show means and s.e.m. across participants. Panels A and B show results for vPulv and LGN respectively. Dashed gray lines show the average monocular responses before and after deprivation. The text inset reports the p-value of the ANOVA time by eye interaction term. **C)** Correlation between deprivation indices computed, for each participant, for BOLD responses in vPulv and for perceptual responses during binocular rivalry (same equation as in Binda et al., 2018); the text inset shows Pearson’s correlation coefficient, and the asterisk marks significance at p < 0.05.

Together, these results suggest that the plasticity effect in the visual thalamus was selective for the vPulv region, where it predicted the perceptual outcome of monocular deprivation.

We performed several control analyses to support these conclusions.

First, we confirmed all our results using two alternative analyses approaches of the fMRI timeseries: General Linear Model (GLM) and Event Related Averaging. Both these methods require assumptions on the temporal dynamics of the BOLD response. GLM relies on choosing an appropriate hemodynamic response function. Using the canonical HRF previously applied to BOLD data from subcortical regions (Koizumi et al., 2019; McFadyen et al., 2019), we confirmed a reliable time by eye interaction in vPulv (F(1,17) = 11.07, p = 0.004), not in LGN (F(1,17) = 0.05, p = 0.822), the two being significantly different as testified by the significant three-way time by eye by ROI interaction (F(1,17) = 6.89, p = 0.018); we also confirmed that the vPulv effect correlated with the behavioral deprivation index (*r*(*18*) = 0.51, p = 0.031). Results were again similar when we quantified BOLD response amplitude from the event-related average curve, which we averaged in the interval between 3s and 12s after using the 0s timepoint for baseline correction (essentially: integrating the response in **Figure 1C** over the 3-12s interval and dividing by the duration of this interval). Again we found a reliable time by eye interaction in vPulv (F(1,17) = 16.07, p = 0.001), not in LGN (F(1,17) = 0.00, p = 0.964), with a significant three-way interaction (F(1,17) = 4.52, p = 0.048) and a significant correlation between the vPulv effect and the behavioral deprivation index (*r*(*18*) = 0.49, p = 0.039).

Second, we checked that our results were not dependent upon the specific definitions of LGN and vPulv regions that we elected to use. To this end, we re-defined ROIs based on two different anatomical templates, intersected with functional activations from a separate dataset. We defined an alternative Pulvinar ROI based on Najdenovska et al.’s atlas (Najdenovska et al., 2018), which was obtained from diffusion-weighted imaging; this label does not separate visual and non-visual subregions of the pulvinar and we used data from an independent experiment involving a subset of our participants (N=9) to isolate the visually responsive subregion. Selecting the 200 most active 1mm^3^ voxels from Najdenovska et al.’s pulvinar, we identified a ventral cluster that largely overlapped the vPulv region used for our main analyses (Allen et al., 2021; Arcaro et al., 2015; Guest et al., 2021) further validating it (**Figure 1-supplement 1**).

Using this alternative definition of vPulv, we still found a significant time by eye interaction (F(1,17) = 13.34, p = 0.002, **Figure 2-supplement 1, panel A**), confirming the reliable monocular deprivation effect in the ventral (or visual) Pulvinar. We followed a similar strategy to obtain an alternative definition of LGN. We located it based on the histological FSL atlas (Burgel et al., 2006; Burgel et al., 1999) and then again analyzed the 200 most active 1mm^3^ voxels (**Figure 1-supplement 1**), thereby equating ROI size between LGN and vPulv. With this alternative definition of LGN, we still found no significant time x eye interaction in LGN (F(1,17) = 0.10 p = 0.756, **Figure 2-supplement 1, panel B**).

## DISCUSSION

Our study is the first to show evidence for short-term plasticity in the adult human thalamus. We found that 2h of monocular deprivation, besides shifting ocular dominance as assessed from behavior and from V1 responses (Binda et al., 2018), also affects ocular drive in a specific subregion of the visual thalamus, ventral Pulvinar or vPulv.

With a series of controls, we obtained strong evidence against the possibility that this is an artifact of BOLD analyses or region labelling; we confirmed the results with three different approaches – we cross-checked them with two independent atlases and reached the same conclusion, that the plasticity effect was clearly observed in vPulv.

In contrast, the adjacent LGN region was reliably unaffected by monocular deprivation. Note that we did not attempt to separate magnocellular and parvocellular subdivisions of the LGN. Previous evidence suggests that this separation is possible with high resolution fMRI (Denison et al., 2014; Qian et al., 2020; Zhang et al., 2015) and indicates that short-term plasticity effect may be strongest for the parvocellular system (Begum & Tso, 2016; Lunghi et al., 2013). However, since parvocellular layers cover the largest majority of the LGN volume, this system should dominate BOLD signals from the LGN region; nevertheless, we found no systematic BOLD modulations with monocular deprivation. Of course, this does not imply that LGN lacks plasticity potential, which may well emerge in other contexts or participants (Castaldi et al., 2016; Jaepel et al., 2017; Levine et al., 2020; Mikellidou et al., 2019; Rose & Bonhoeffer, 2018) or during a stabilization of the short-term plasticity effect, as observed for repeated monocular deprivations in amblyopia (Lunghi et al., 2019). While we cannot exclude that plasticity is possible in LGN, the fact that vPulv responses clearly changed after monocular deprivation in the same dataset where LGN showed no modulation suggests that short-term plasticity effects in LGN, if present, are small or inconsistent.

With BOLD data, it is not possible to ascertain whether the plasticity effect seen in the thalamus is a feed-forward or feed-back effect, i.e. whether it is generated intra-cortically and merely extended to the thalamus due to the tight cortico-thalamic feedback connections, or whether the plasticity effect originates within the thalamus, which projects it to the cortex via feedforward connections.

The first hypothesis is consistent with evidence that ocular dominance plasticity (both long- and short-term) depends on features and processes that are intrinsic to the primary visual cortex, such as intracortical inhibition (Fagiolini & Hensch, 2000; Heimel et al., 2011; Hensch et al., 1998; Lunghi, Emir, et al., 2015; Sale et al., 2010). It also fits with evidence that feedback connections are more susceptible to homeostatic plasticity than feedforward thalamo-cortical connections (Krupa et al., 1999; Miska et al., 2018). Moreover, the distribution of feedback signals from the visual cortex to LGN and vPulv may help explain the difference in behavior of these thalamic regions. While LGN bidirectional connections are mainly with V1 (Briggs & Usrey, 2011), vPulv serves as a hub for converging feedback from vast portions of the occipital and temporal cortex (Arcaro et al., 2018). There is evidence that vPulv primarily connects with ventral visual cortical areas (Arcaro et al., 2018) – although connections with MT were also documented, especially early in development (Bridge et al., 2016). Our previous work indicates that short-term plasticity effects extend beyond primary visual cortex mainly to the ventral visual stream (Binda et al., 2018). Thus, vPulv is broadly connected with a large cortical territory where short-term plasticity is the strongest; if short-term plasticity effects are carried through feedback signals, it is reasonable to assume that these will be stronger, more stable, and ultimately easier to detect in vPulv than in LGN.

The second hypothesis, that plasticity effects are (at least in part) generated within the thalamus, is in line with growing evidence on the importance of the thalamus in active vision (Saalmann & Kastner, 2011). The traditional view of this nucleus as a passive relay of peripheral information has been overruled by evidence that the thalamus actively regulates information transmission to the cortex and between cortical areas (Saalmann & Kastner, 2011). This is particularly true for the pulvinar, which has been involved in a variety of mechanisms, including the modulation of response magnitude through gain control (Fiebelkorn & Kastner, 2019; Purushothaman et al., 2012) and synchrony of neurons (Saalmann et al., 2012) according to behavioral demands. These may be implemented through loops of cortico-pulvinar-cortical pathways (Jaramillo et al., 2019), which allow for filtering or gating incoming information. These functions have been often studied in the context of attention (Zhou et al., 2016); however, gain control of cortical responses is likely to participate in setting ocular dominance and regulating its short-term changes (Lunghi et al., 2011; Spiegel et al., 2017).

Although these two hypotheses (that short-term plasticity affects feedback or feedforward connections between the thalamus and the visual cortex) are equally compatible with the bulk of our data, the concept of plasticity originating in the pulvinar may be better suited to explain the correlation between BOLD modulations in this subcortical area and perceptual modulations. Interestingly, the concept that the visual pulvinar plays a fundamental role in short-term plasticity is also supported by a recent human neuroimaging study, where pulvinar was suggested to gate GABAergic inhibition in the cortex and the associated short-term learning effect (Ziminski et al., 2021).

In conclusion, the present study showed that short-term monocular deprivation effects, which are widespread in cortical visual areas, also extend to subcortical regions. Within the thalamus, plasticity mainly affects the ventral portion of the pulvinar – the portion of this nucleus that shows the most obvious visual functions and the strongest recurrent connections with visual cortex. Although our fMRI data do not allow to ascertain the origin of the plasticity effect, our observations open the possibility that ventral pulvinar plays a role in setting ocular dominance and maintaining its plasticity in adulthood.

## METHODS

### Participants and Monocular Deprivation procedure

Experimental procedures are in line with the declaration of Helsinki and were approved by the regional ethics committee [Comitato Etico Pediatrico Regionale—Azienda Ospedaliero-Universitaria Meyer—Firenze (FI)] and by the Italian Ministry of Health, under the protocol ‘Plasticità e multimodalità delle prime aree visive: studio in risonanza magnetica a campo ultra alto (7T)’ version #1 dated 11/11/2015. Written informed consent was obtained from each participant, which included consent to process and preserve the data and publish them in anonymous form. Twenty healthy volunteers with normal or corrected-to-normal visual acuity were examined (8 females and 12 males, mean age = 27 years). Sample size was set based on the minimum number of participants (N = 17) required to reliably detect a medium sized correlation between MRI and psychophysical data: r = 0.62 or higher, as reported in previous MR work on short-term plasticity (Lunghi, Emir, et al., 2015). Two (male) participants were excluded. One because of strong eye dominance (already excluded for the analyses in Binda et al., 2018) and the second due to a large vein passing near LGN that could bias the BOLD response. This left 18 participants (8 females and 10 males). We analyzed data from two fMRI sessions, before and after 2h of monocular deprivation, performed by patching the dominant eye with a translucent patch. Binocular rivalry was measured immediately before each fMRI session to estimate the transient ocular dominance shift. The effect of deprivation on perception and brain activations was estimated by computing a deprivation index (Binda et al., 2018). This is the post-to pre-deprivation ratio of values “y” for the deprived eye, divided by the same value for the non-deprived eye:

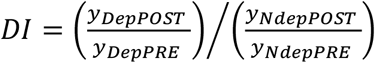

with “y” defined as binocular rivalry phase durations or BOLD responses. Using the same equation to compute the deprivation effects on psychophysical and BOLD data allowed for correlating them across participants (Figure 2C).

### fMRI acquisition protocols and analyses

Detailed information on the protocol and data preprocessing may be found in Binda et al. (2018). In our previous publication, we limited our analyses to the cortical projections of fMRI time series and focused on BOLD responses in the visual cortex. Here we analyzed fMRI time series in the volume and focused on subcortical visual structures. Individual participants’ data were aligned to a standard anatomical template, the MNI atlas, using ANTs (Avants et al., 2008; Avants et al., 2011). ANTs aligned T1 anatomical images (acquired with 1 mm isotropic resolution) to the MNI template available in FSL (Collins et al., 1995; Mazziotta et al., 2001), by means of an affine registration matrix and a warp field. These were used to transform individual participants’ preprocessed BOLD data (EPI-GRE with 1.5 mm isotropic resolution and TR = 3s, which had been slice-time, motion and distortion corrected) to the MNI space through the antsRegistrationSyN.sh routine (Tustison & Avants, 2013).

BOLD time series were averaged across voxels within each ROI (see below), resulting in one time series per each of the 18 participants, two ROIs and four conditions (stimulating the deprived and non-deprived eye, before and after monocular deprivation). Individual BOLD time series were transformed into percent signal change units (by subtracting and dividing by the mean signal) and detrended.

We acquired four BOLD time series per participant, two before and two after monocular deprivation. In each series, only one eye was stimulated, and the other viewed a mid-level gray image. Stimuli consisted of bandpass filtered, dynamic noise images presented in a block design, with 9 sec long periods of stimulation (during which the noise stimulus was refreshed at a rate of 8 Hz) separated by 12 sec of rest (mid-level gray screen), repeated 10 times. Across blocks, the spatial frequency cut-off of the bandpass filter was varied. Unlike in Binda et al. (2018), here we pooled across spatial frequencies, for both theorical (spatial frequency tuning in the thalamus is not expected to be as sharp as in the cortex) and practical reasons (pooling across repetitions compensates for the lower SNR of the subcortical regions). This turned our stimulus into a periodic alternation of ON (9 sec) and OFF periods (12 sec), expected to generate periodic visually evoked responses, the amplitude of which can be efficiently estimated with Fourier analysis, extracting the amplitude at the stimulus frequency (1 cycle every 21 sec or 0.047 Hz). The advantage of this method is that it does not make assumptions on the latency of the response, which is captured by the phase parameter, and it is free to vary across regions. For visualization purposes (Figure 1A and Figure 1-supplement 1) amplitude estimates were computed for individual voxels, after averaging fMRI time series across conditions and participants (amplitude estimates were divided by the root mean squared error of the corresponding sinusoidal function, yielding a value that is conceptually similar to a t-statistics).

We complemented this analysis with two other methods that, contrary to the Fourier approach, do make assumptions on response latencies.

First, we used an Event related averaging approach to estimate the profiles of fMRI responses. We selected 21 sec long (7TRs) BOLD epochs following each stimulus onset and averaged across epochs (of which we had 10 per acquisition). We assumed that the response occurs between 3 sec and 12 sec from stimulus onset, and we used the average over this interval to estimate its amplitude.

Second, we used a General Linear Modelling, and we assumed a canonical (two-gamma) hemodynamic response function (HRF) as previously used to model subcortical responses (Koizumi et al., 2019; McFadyen et al., 2019)

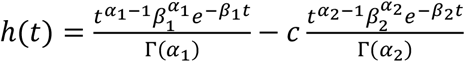

where *t* is time, α_1_ = 6, α_2_ = 16, β_1_ = β_2_ = 1, *c* = 1/6 and Γ represents the gamma function.

We generated a stimulus predictor (boxcar function representing the stimulus ON/OFF periods, convolved by the HRF) and two nuisance predictors (a linear trend and a constant) and we extracted the corresponding beta-weights by linear regression.

### ROI definition

Thalamic ROIs were defined in the MNI space based on publicly available atlases.

ROIs for the main analysis were taken from the recently published NSD dataset, for which they were defined based on a combination of functional data (retinotopic mapping experiments) constrained with anatomical features (Allen et al., 2021; Arcaro et al., 2015).

ROIs for the confirmatory analyses were based on two additional MNI atlases. Pulvinar was labelled according to an atlas developed from diffusion tensor imaging data (Najdenovska et al., 2018); LGN was labelled according to an histological atlas available in FSL (Burgel et al., 2006; Burgel et al., 1999), setting coverage threshold to 50%. These anatomical labels were intersected with a map of visual responses, which we measured in a separate experiment with a subset of our participants (N = 9). This consisted of four 70 TR long series, presenting full contrast full screen binary noise images (random checks of variable size dynamically refreshed at 8 Hz) for epochs of 9 s (3 TRs) separated by 12 sec epochs of mid-level gray (4 TRs). Visual activations were measured with the Fourier approach. In each hemisphere and ROI, we selected the 200 most active voxels (highest activity), and we used these to define two visually responsive ROIs of equal size within the anatomically defined LGN and Pulvinar regions.

---

**Figure 1-supplement 1.**
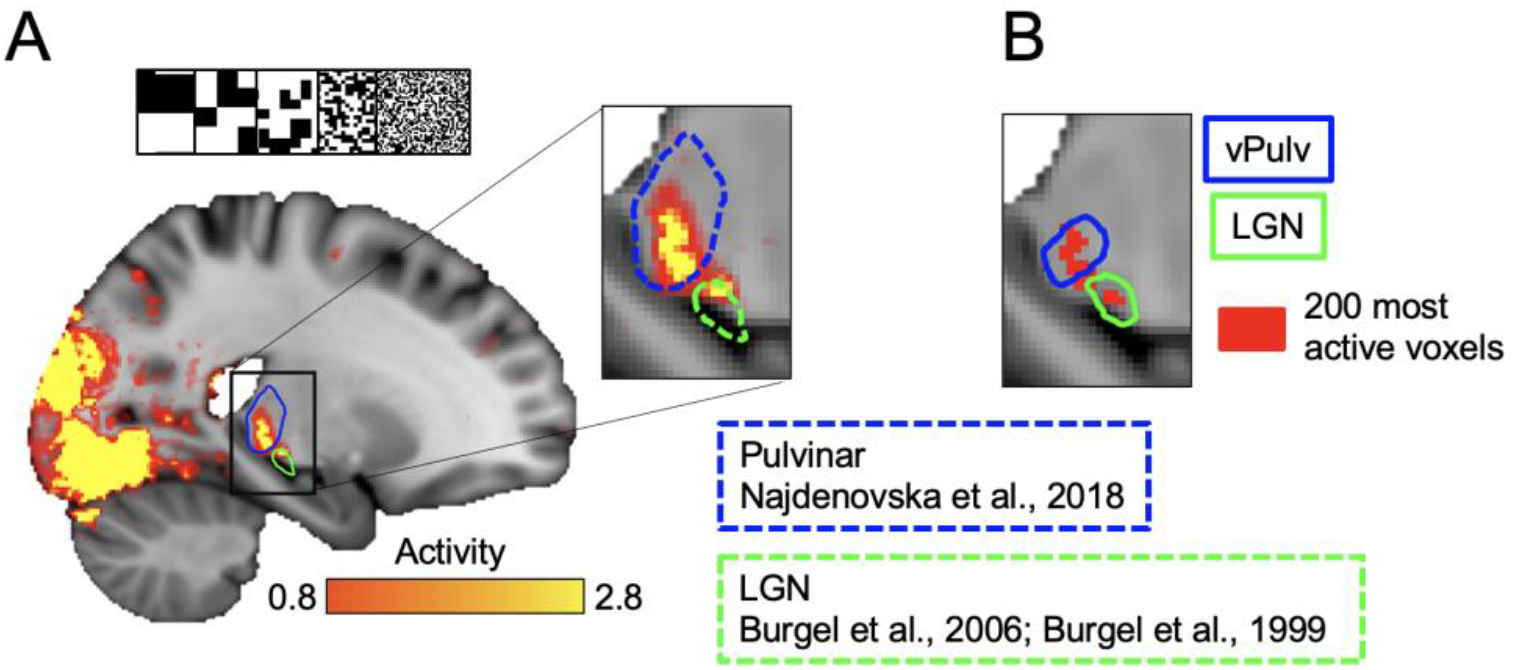
Alternative definition of thalamic ROIs. **A)** Map of activity evoked by another visual stimulus (binary noise, shown at the top, tested in a subset of N=9 participants), estimated by fast Fourier analysis of the fMRI timeseries. Dashed blue and green outlines show the anatomical templates (Burgel et al., 2006; Burgel et al., 1999; Najdenovska et al., 2018) that we used in conjunction with the functional activations (the 200 most active voxels within each anatomical mask, shown in panel B) to define alternative ROIs for visual Pulvinar and LGN. **B)** Most active voxels within the anatomical templates in A, compared with the outline of ROIs used for the main analyses (Allen et al., 2021).

**Figure 2-supplement 1.**
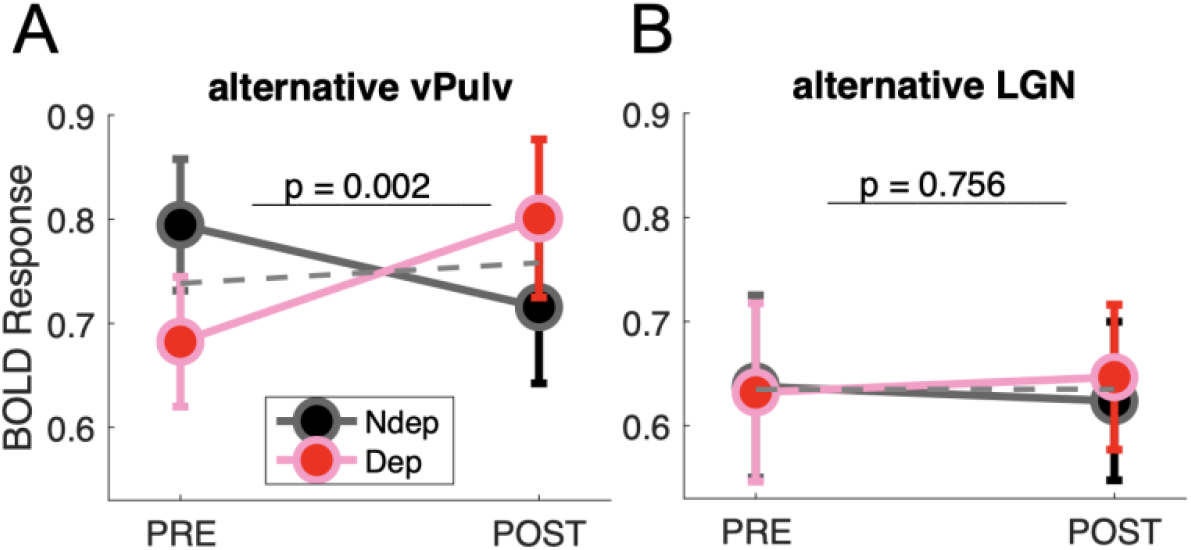
Alternative ROI definition confirms short-term plasticity in Pulvinar, not in LGN. Modulation of BOLD responses with monocular deprivation in visual Pulvinar (**A**) and LGN (**B**), as defined by anatomically defined ROIs (Burgel et al., 2006; Burgel et al., 1999; Najdenovska et al., 2018) intersected with functional activations (Figure 1-supplement 1). Same conventions as in Figure 2.

## Acknowledgments

This work was supported by the European Research Council (ERC) under the European Union’s Horizon 2020 research and innovation program, grant n. 801715 (PUPILTRAITS) and n. 832813 (GenPercept), and by the Italian Ministry of University and Research under the PRIN2017 programme (grant MISMATCH) and FARE-2 (grant SMILY).

## References

Allen, E. J., St-Yves, G., Wu, Y., Breedlove, J. L., Dowdle, L. T., Caron, B., Pestilli, F., Charest, I., Hutchinson, J. B., Naselaris, T., & Kay, K. (2021). A massive 7T fMRI dataset to bridge cognitive and computational neuroscience. bioRxiv, 2021.2002.2022.432340. https://doi.org/10.1101/2021.02.22.432340

Arcaro, M. J., Pinsk, M. A., Chen, J., & Kastner, S. (2018, Dec 19). Organizing principles of pulvino-cortical functional coupling in humans. Nat Commun, 9(1), 5382. https://doi.org/10.1038/s41467-018-07725-6

Arcaro, M. J., Pinsk, M. A., & Kastner, S. (2015, Jul 8). The Anatomical and Functional Organization of the Human Visual Pulvinar. J Neurosci, 35(27), 9848–9871. https://doi.org/10.1523/JNEUROSCI.1575-14.2015

Avants, B. B., Epstein, C. L., Grossman, M., & Gee, J. C. (2008, Feb). Symmetric diffeomorphic image registration with cross-correlation: evaluating automated labeling of elderly and neurodegenerative brain. Med Image Anal, 12(1), 26–41. https://doi.org/10.1016/j.media.2007.06.004

Avants, B. B., Tustison, N. J., Song, G., Cook, P. A., Klein, A., & Gee, J. C. (2011, Feb 1). A reproducible evaluation of ANTs similarity metric performance in brain image registration. Neuroimage, 54(3), 2033–2044. https://doi.org/10.1016/j.neuroimage.2010.09.025

Bai, J., Dong, X., He, S., & Bao, M. (2017, Jun 3). Monocular deprivation of Fourier phase information boosts the deprived eye’s dominance during interocular competition but not interocular phase combination. Neuroscience, 352, 122–130. https://doi.org/10.1016/j.neuroscience.2017.03.053

Begum, M., & Tso, D. (2016). Shifts in interocular balance resulting from short-term monocular deprivation in adult macaque visual cortex are not magno-dominated. Journal of Vision, 16(12), 1328–1328. https://doi.org/10.1167/16.12.1328

Binda, P., Kurzawski, J. W., Lunghi, C., Biagi, L., Tosetti, M., & Morrone, M. C. (2018, Nov 26). Response to short-term deprivation of the human adult visual cortex measured with 7T BOLD. Elife, 7. https://doi.org/10.7554/eLife.40014

Binda, P., & Lunghi, C. (2017). Short-Term Monocular Deprivation Enhances Physiological Pupillary Oscillations. Neural Plast, 2017, 6724631. https://doi.org/10.1155/2017/6724631

Blasdel, G. G., & Lund, J. S. (1983, Jul). Termination of afferent axons in macaque striate cortex. J Neurosci, 3(7), 1389–1413. https://www.ncbi.nlm.nih.gov/pubmed/6864254

Bourne, J. A., & Morrone, M. C. (2017). Plasticity of Visual Pathways and Function in the Developing Brain: Is the Pulvinar a Crucial Player? Front Syst Neurosci, 11, 3. https://doi.org/10.3389/fnsys.2017.00003

Bridge, H., Leopold, D. A., & Bourne, J. A. (2016, Feb). Adaptive Pulvinar Circuitry Supports Visual Cognition. Trends Cogn Sci, 20(2), 146–157. https://doi.org/10.1016/j.tics.2015.10.003

Briggs, F., & Usrey, W. M. (2011, Jan 1). Corticogeniculate feedback and visual processing in the primate. J Physiol, 589(Pt 1), 33–40. https://doi.org/10.1113/jphysiol.2010.193599

Burgel, U., Amunts, K., Hoemke, L., Mohlberg, H., Gilsbach, J. M., & Zilles, K. (2006, Feb 15). White matter fiber tracts of the human brain: three-dimensional mapping at microscopic resolution, topography and intersubject variability. Neuroimage, 29(4), 1092–1105. https://doi.org/10.1016/j.neuroimage.2005.08.040

Burgel, U., Schormann, T., Schleicher, A., & Zilles, K. (1999, Nov). Mapping of histologically identified long fiber tracts in human cerebral hemispheres to the MRI volume of a reference brain: position and spatial variability of the optic radiation. Neuroimage, 10(5), 489–499. https://doi.org/10.1006/nimg.1999.0497

Castaldi, E., Cicchini, G. M., Cinelli, L., Biagi, L., Rizzo, S., & Morrone, M. C. (2016, Oct). Visual BOLD Response in Late Blind Subjects with Argus II Retinal Prosthesis. PLoS Biol, 14(10), e1002569. https://doi.org/10.1371/journal.pbio.1002569

Castaldi, E., Lunghi, C., & Morrone, M. C. (2020, May). Neuroplasticity in adult human visual cortex. Neurosci Biobehav Rev, 112, 542–552. https://doi.org/10.1016/j.neubiorev.2020.02.028

Chadnova, E., Reynaud, A., Clavagnier, S., & Hess, R. F. (2017, Feb 2). Short-term monocular occlusion produces changes in ocular dominance by a reciprocal modulation of interocular inhibition. Sci Rep, 7, 41747. https://doi.org/10.1038/srep41747

Collins, D. L., Holmes, C. J., Peters, T. M., & Evans, A. C. (1995, 1995/01/01). Automatic 3-D model-based neuroanatomical segmentation [https://doi.org/10.1002/hbm.460030304]. Human Brain Mapping, 3(3), 190–208. https://doi.org/10.1002/hbm.460030304

Denison, R. N., Vu, A. T., Yacoub, E., Feinberg, D. A., & Silver, M. A. (2014, Nov 15). Functional mapping of the magnocellular and parvocellular subdivisions of human LGN. Neuroimage, 102 Pt 2, 358–369. https://doi.org/10.1016/j.neuroimage.2014.07.019

Fagiolini, M., & Hensch, T. K. (2000, Mar 9). Inhibitory threshold for critical-period activation in primary visual cortex. Nature, 404(6774), 183–186. https://doi.org/10.1038/35004582

Fiebelkorn, I. C., & Kastner, S. (2019, Jan 16). The Puzzling Pulvinar. Neuron, 101(2), 201–203. https://doi.org/10.1016/j.neuron.2018.12.032

Guest, D., Allen, E., Wu, Y., Naselaris, T., Arcaro, M., & Kay, K. (2021). Evidence for a ventral visual stream in the pulvinar. Journal of Vision, 21(9), 2809–2809. https://doi.org/10.1167/jov.21.9.2809

Heimel, J. A., van Versendaal, D., & Levelt, C. N. (2011). The role of GABAergic inhibition in ocular dominance plasticity. Neural Plast, 2011, 391763. https://doi.org/10.1155/2011/391763

Hendrickson, A. E., Wilson, J. R., & Ogren, M. P. (1978, Nov 1). The neuroanatomical organization of pathways between the dorsal lateral geniculate nucleus and visual cortex in Old World and New World primates. J Comp Neurol, 182(1), 123–136. https://doi.org/10.1002/cne.901820108

Hensch, T. K., Fagiolini, M., Mataga, N., Stryker, M. P., Baekkeskov, S., & Kash, S. F. (1998, Nov 20). Local GABA circuit control of experience-dependent plasticity in developing visual cortex. Science, 282(5393), 1504–1508. https://doi.org/10.1126/science.282.5393.1504

Hensch, T. K., & Quinlan, E. M. (2018, Jan). Critical periods in amblyopia. Vis Neurosci, 35, E014. https://doi.org/10.1017/S0952523817000219

Hubel, D. H., & Wiesel, T. N. (1972, Dec). Laminar and columnar distribution of geniculo-cortical fibers in the macaque monkey. J Comp Neurol, 146(4), 421–450. https://doi.org/10.1002/cne.901460402

Jaepel, J., Hubener, M., Bonhoeffer, T., & Rose, T. (2017, Dec). Lateral geniculate neurons projecting to primary visual cortex show ocular dominance plasticity in adult mice. Nat Neurosci, 20(12), 1708–1714. https://doi.org/10.1038/s41593-017-0021-0

Jaramillo, J., Mejias, J. F., & Wang, X. J. (2019, Jan 16). Engagement of Pulvino-cortical Feedforward and Feedback Pathways in Cognitive Computations. Neuron, 101(2), 321–336 e329. https://doi.org/10.1016/j.neuron.2018.11.023

Koizumi, A., Zhan, M., Ban, H., Kida, I., De Martino, F., Vaessen, M. J., de Gelder, B., & Amano, K. (2019, Nov/Dec). Threat Anticipation in Pulvinar and in Superficial Layers of Primary Visual Cortex (V1). Evidence from Layer-Specific Ultra-High Field 7T fMRI. eNeuro, 6(6). https://doi.org/10.1523/ENEURO.0429-19.2019

Krupa, D. J., Ghazanfar, A. A., & Nicolelis, M. A. (1999, Jul 6). Immediate thalamic sensory plasticity depends on corticothalamic feedback. Proc Natl Acad Sci U S A, 96(14), 8200–8205. https://doi.org/10.1073/pnas.96.14.8200

Levine, A. T., Yuen, K., Gouws, A., Wade, A. R., Morland, A. B., Codina, C., Buckley, D., & Baseler, H. A. (2020). Retinotopic remapping of the visual system in deaf adults. bioRxiv, 2020.2001.2031.923342. https://doi.org/10.1101/2020.01.31.923342

Lewis, L. D., Setsompop, K., Rosen, B. R., & Polimeni, J. R. (2018, Nov 1). Stimulus-dependent hemodynamic response timing across the human subcortical-cortical visual pathway identified through high spatiotemporal resolution 7T fMRI. Neuroimage, 181, 279–291. https://doi.org/10.1016/j.neuroimage.2018.06.056

Lunghi, C., Berchicci, M., Morrone, M. C., & Di Russo, F. (2015, Oct 1). Short-term monocular deprivation alters early components of visual evoked potentials. J Physiol, 593(19), 4361–4372. https://doi.org/10.1113/JP270950

Lunghi, C., Burr, D. C., & Morrone, C. (2011, Jul 26). Brief periods of monocular deprivation disrupt ocular balance in human adult visual cortex. Curr Biol, 21(14), R538–539. https://doi.org/10.1016/j.cub.2011.06.004

Lunghi, C., Burr, D. C., & Morrone, M. C. (2013, May 1). Long-term effects of monocular deprivation revealed with binocular rivalry gratings modulated in luminance and in color. J Vis, 13(6). https://doi.org/10.1167/13.6.1

Lunghi, C., Emir, U. E., Morrone, M. C., & Bridge, H. (2015, Jun 1). Short-term monocular deprivation alters GABA in the adult human visual cortex. Curr Biol, 25(11), 1496–1501. https://doi.org/10.1016/j.cub.2015.04.021

Lunghi, C., & Sale, A. (2015, Dec 7). A cycling lane for brain rewiring. Curr Biol, 25(23), R1122–1123. https://doi.org/10.1016/j.cub.2015.10.026

Lunghi, C., Sframeli, A. T., Lepri, A., Lepri, M., Lisi, D., Sale, A., & Morrone, M. C. (2019, Feb). A new counterintuitive training for adult amblyopia. Ann Clin Transl Neurol, 6(2), 274–284. https://doi.org/10.1002/acn3.698

Lyu, L., He, S., Jiang, Y., Engel, S. A., & Bao, M. (2020, May 21). Natural-scene-based Steady-state Visual Evoked Potentials Reveal Effects of Short-term Monocular Deprivation. Neuroscience, 435, 10–21. https://doi.org/10.1016/j.neuroscience.2020.03.039

Mazziotta, J., Toga, A., Evans, A., Fox, P., Lancaster, J., Zilles, K., Woods, R., Paus, T., Simpson, G., Pike, B., Holmes, C., Collins, L., Thompson, P., MacDonald, D., Iacoboni, M., Schormann, T., Amunts, K., Palomero-Gallagher, N., Geyer, S., Parsons, L., Narr, K., Kabani, N., Le Goualher, G., Boomsma, D., Cannon, T., Kawashima, R., & Mazoyer, B. (2001, Aug 29). A probabilistic atlas and reference system for the human brain: International Consortium for Brain Mapping (ICBM). Philos Trans R Soc Lond B Biol Sci, 356(1412), 1293–1322. https://doi.org/10.1098/rstb.2001.0915

McFadyen, J., Mattingley, J. B., & Garrido, M. I. (2019, Jan 16). An afferent white matter pathway from the pulvinar to the amygdala facilitates fear recognition. Elife, 8. https://doi.org/10.7554/eLife.40766

Mikellidou, K., Arrighi, R., Aghakhanyan, G., Tinelli, F., Frijia, F., Crespi, S., De Masi, F., Montanaro, D., & Morrone, M. C. (2019, May). Plasticity of the human visual brain after an early cortical lesion. Neuropsychologia, 128, 166–177. https://doi.org/10.1016/j.neuropsychologia.2017.10.033

Min, S. H., Baldwin, A. S., Reynaud, A., & Hess, R. F. (2018, Nov 20). The shift in ocular dominance from short-term monocular deprivation exhibits no dependence on duration of deprivation. Sci Rep, 8(1), 17083. https://doi.org/10.1038/s41598-018-35084-1

Miska, N. J., Richter, L. M., Cary, B. A., Gjorgjieva, J., & Turrigiano, G. G. (2018, Oct 12). Sensory experience inversely regulates feedforward and feedback excitation-inhibition ratio in rodent visual cortex. Elife, 7. https://doi.org/10.7554/eLife.38846

Najdenovska, E., Aleman-Gomez, Y., Battistella, G., Descoteaux, M., Hagmann, P., Jacquemont, S., Maeder, P., Thiran, J. P., Fornari, E., & Bach Cuadra, M. (2018, Nov 27). In-vivo probabilistic atlas of human thalamic nuclei based on diffusion-weighted magnetic resonance imaging. Sci Data, 5, 180270. https://doi.org/10.1038/sdata.2018.270

Purushothaman, G., Marion, R., Li, K., & Casagrande, V. A. (2012, Jun). Gating and control of primary visual cortex by pulvinar. Nat Neurosci, 15(6), 905–912. https://doi.org/10.1038/nn.3106

Qian, Y., Zou, J., Zhang, Z., An, J., Zuo, Z., Zhuo, Y., Wang, D. J. J., & Zhang, P. (2020, Apr 29). Robust functional mapping of layer-selective responses in human lateral geniculate nucleus with high-resolution 7T fMRI. Proc Biol Sci, 287(1925), 20200245. https://doi.org/10.1098/rspb.2020.0245

Rose, T., & Bonhoeffer, T. (2018, Dec). Experience-dependent plasticity in the lateral geniculate nucleus. Curr Opin Neurobiol, 53, 22–28. https://doi.org/10.1016/j.conb.2018.04.016

Saalmann, Y. B., & Kastner, S. (2011, Jul 28). Cognitive and perceptual functions of the visual thalamus. Neuron, 71(2), 209–223. https://doi.org/10.1016/j.neuron.2011.06.027

Saalmann, Y. B., Pinsk, M. A., Wang, L., Li, X., & Kastner, S. (2012, Aug 10). The pulvinar regulates information transmission between cortical areas based on attention demands. Science, 337(6095), 753–756. https://doi.org/10.1126/science.1223082

Sale, A., Berardi, N., Spolidoro, M., Baroncelli, L., & Maffei, L. (2010). GABAergic inhibition in visual cortical plasticity. Front Cell Neurosci, 4, 10. https://doi.org/10.3389/fncel.2010.00010

Schwenk, J. C. B., VanRullen, R., & Bremmer, F. (2020). Dynamics of Visual Perceptual Echoes Following Short-Term Visual Deprivation. Cereb Cortex Commun, 1(1), tgaa012. https://doi.org/10.1093/texcom/tgaa012

Sommeijer, J. P., Ahmadlou, M., Saiepour, M. H., Seignette, K., Min, R., Heimel, J. A., & Levelt, C. N. (2017, Dec). Thalamic inhibition regulates critical-period plasticity in visual cortex and thalamus. Nat Neurosci, 20(12), 1715–1721. https://doi.org/10.1038/s41593-017-0002-3

Spiegel, D. P., Baldwin, A. S., & Hess, R. F. (2017, Jan 10). Ocular dominance plasticity: inhibitory interactions and contrast equivalence. Sci Rep, 7, 39913. https://doi.org/10.1038/srep39913

Turrigiano, G. (2012, Jan 1). Homeostatic synaptic plasticity: local and global mechanisms for stabilizing neuronal function. Cold Spring Harb Perspect Biol, 4(1), a005736. https://doi.org/10.1101/cshperspect.a005736

Tustison, N. J., & Avants, B. B. (2013). Explicit B-spline regularization in diffeomorphic image registration. Front Neuroinform, 7, 39. https://doi.org/10.3389/fninf.2013.00039

Wang, M., McGraw, P., & Ledgeway, T. (2020, Aug). Short-term monocular deprivation reduces inter-ocular suppression of the deprived eye. Vision Res, 173, 29–40. https://doi.org/10.1016/j.visres.2020.05.001

Wiesel, T. N., & Hubel, D. H. (1963, Nov). Effects of Visual Deprivation on Morphology and Physiology of Cells in the Cats Lateral Geniculate Body. J Neurophysiol, 26, 978–993. https://doi.org/10.1152/jn.1963.26.6.978

Wiesel, T. N., & Hubel, D. H. (1965, Nov). Comparison of the effects of unilateral and bilateral eye closure on cortical unit responses in kittens. J Neurophysiol, 28(6), 1029–1040. https://doi.org/10.1152/jn.1965.28.6.1029

Zhang, P., Zhou, H., Wen, W., & He, S. (2015, May 1). Layer-specific response properties of the human lateral geniculate nucleus and superior colliculus. Neuroimage, 111, 159–166. https://doi.org/10.1016/j.neuroimage.2015.02.025

Zhou, H., Schafer, R. J., & Desimone, R. (2016, Jan 6). Pulvinar-Cortex Interactions in Vision and Attention. Neuron, 89(1), 209–220. https://doi.org/10.1016/j.neuron.2015.11.034

Zhou, J., Baker, D. H., Simard, M., Saint-Amour, D., & Hess, R. F. (2015). Short-term monocular patching boosts the patched eye’s response in visual cortex. Restor Neurol Neurosci, 33(3), 381–387. https://doi.org/10.3233/RNN-140472

Zhou, J., Clavagnier, S., & Hess, R. F. (2013, Apr 18). Short-term monocular deprivation strengthens the patched eye’s contribution to binocular combination. J Vis, 13(5). https://doi.org/10.1167/13.5.12

Zhou, J., Reynaud, A., & Hess, R. F. (2014, Nov 22). Real-time modulation of perceptual eye dominance in humans. Proc Biol Sci, 281(1795). https://doi.org/10.1098/rspb.2014.1717

Ziminski, J. J., Frangou, P., Karlaftis, V. M., & Kourtzi, Z. (2021). Pulvinar plasticity gates inhibitory processing in visual cortex for perceptual learning. Meeting of the Organization for Human Brain Mapping (2021), https://ww4.aievolution.com/hbm2101/index.cfm?do=abs.viewAbs&src=ext&abs=1838

